# Neural mechanisms underlying anxiety-related deficits of attentional inhibition: direct ERP evidence from the Pd component

**DOI:** 10.1101/2021.10.09.463788

**Authors:** Hu Liping, Huikang Zhang, Tang Hongsi, Lin Shen, Rui Wu, Huang Yan

## Abstract

Behavioral evidence shows that anxious individuals tend to be distracted by irrelevant stimulation not only for threat-related stimuli but also for non-emotional neutral stimuli. These findings suggest that anxious individuals may have a general impairment of attentional control, especially inhibition function. However, the neural mechanism underlying the anxiety-related impairment in attentional control is unclear. Here, in a visual search task with geometric stimuli, we examined attentional processing of the non-emotional neutral distractor on participants with different levels of anxiety, using the event-related-potential (ERP) indices of attentional selection (N2 posterior contralateral [N2pc]) and top-down inhibition (distractor positivity [Pd]). We found that distractor-evoked Pd amplitudes were negatively correlated with trait-anxiety scores, i.e., the higher the level of anxiety, the worse the ability of attentional inhibition. In contrast, the amplitudes of distractor-evoked N2pc did not vary with anxiety levels, suggesting that trait-anxiety level does not affect stimulus-driven attentional capture. We also observed attentional processing of target stimuli and found that the peak latency of target-evoked N2pc was delayed as anxiety levels rise, suggesting that anxiety impairs the efficiency of top-down attentional selection of the target. The present study provides direct neurophysiological evidence for general anxiety-related impairment of attentional control.

## 1. Introduction

Trait anxiety describes the relatively stable and enduring tendency to experience anxiety. Long-term anxiety is reported to impair cognitive function, such as attentional control (Wu & Yan, 2017; Osinsky et al., 2012). Previous studies have shown that high-level trait anxiety is associated with strong interference from task-irrelevant threat distractors, such as angry and fearful faces, or threat-related words (Bar-Haim et al., 2007; Dennis & Chen, 2009; McTeague et al., 2011; Gutierrez & Berggren, 2020). Recent research shows that a non-emotional neutral distractor also triggers stronger interference for individuals with a higher level of anxiety (Berggren & Derakshan, 2014; Moran & Moser, 2015; Moser et al., 2012; Pacheco-Ungietti et al., 2010). These findings suggest that trait anxiety is associated with more general attentional and cognitive control deficits, not just a bias toward negative emotional stimuli.

Most cognitive models of anxiety agree that anxiety disrupts the balance between the stimulus-driven/bottom-up attention system and the goal-directed/top-down attention system. For example, attentional control theory puts forward that anxiety increases the bottom-up attention and reduces the top-down attentional control, especially inhibition function (Eysenck et al., 2007; Eysenck & Derakshan, 2011). A similar perspective was proposed by Mogg and Bradley (2018) in their cognitive-motivational framework. Some behavioral evidence supports that an attentional bias (stimulus-driven attention) towards the salient distractors may contribute to the stronger interference effect in anxious individuals (Bar-Haim et al., 2007; Moser et al., 2012; Gutierrez & Berggren, 2020). An electroencephalography (EEG) study provides neural evidence that the non-emotional salient distractor captures more attention of highly anxious individuals, evoking a larger distractor-evoked N2pc component (contralateral-minus-ipsilateral negative potential, an indicator of attentional selection, Gasper et al, 2018). On the other hand, anxiety may be associated with reduced top-down attentional control, in which the inhibitory system plays a crucial part by filtering task-irrelevant information (Gaspelin & Luck, 2017). Some behavioral findings suggest that impairment in inhibitory control may account for the larger interference effect associated with high anxiety (Berggren & Derakshan, 2014; Kalanthroff et al., 2016; Wieser et al., 2009). Yet, relatively few studies report neural mechanisms underlying the anxiety-related deficit of top-down inhibition. Some functional magnetic resonance imaging (fMRI) evidence suggests that reduced prefrontal attentional control is linked to impaired inhibitory control in highly anxious individuals (Basten et al., 2011; Bishop, 2009). In EEG studies, the Pd component (contralateral-minus-ipsilateral positive potential) is generally accepted as the direct neural indicator of top-down attentional inhibition (Gaspelin & Luck, 2017; Sawaki & Luck, 2010; Wang et al., 2016). However, so far no studies have found direct neurophysiological evidence to support anxiety-related impairment of attentional inhibition from the Pd, which is a direct indicator of attentional inhibition.

The present study aims to investigate the neural mechanism underlying the anxiety-related deficits of attentional control in processing a general (i.e., non-emotion-specific) visual stimulus. We examine attentional processing through specific ERP components, i.e., the N2pc (attentional selection) and the Pd (attentional inhibition) components (Luck & Hillyard, 1994; Hickey et al., 2009). We use a visual search paradigm with a task-irrelevant color-singleton distractor and a changeable target. Specifically, the target was designated as a unique geometrical shape among the six items, i.e., searching for a diamond in circles or searching for a circle in diamonds. The shape of target and the color of distractor were randomly switched between trials. The color distractor may automatically attract spatial attention (evoking the N2pc), or be suppressed by top-down modulation (evoking the Pd). We examine which process accounts for the interference effect associated with anxiety: if the distractor-evoked N2pc amplitudes increase as anxiety levels rise, it means that anxiety elicits more attentional capture towards the distractor (i.e., bottom-up attention increases); or if the Pd amplitudes reduce as anxiety levels rise, it then means that anxiety impairs the inhibition function (i.e., top-down inhibition decreases). Additionally, we also examine the target-evoked N2pc to investigate whether top-down attentional selection of the target is affected by anxiety levels. To observe the correlation between ERP indicators of attentional processing and trait-anxiety level, we recruited participants with different levels of trait anxiety.

## 2. Materials and methods

### 2.1 Ethics statement

The present study was conducted following the tenets of the World Medical Association Declaration of Helsinki and was approved by the Human Research Ethics Committee of the Shenzhen Institutes of Advanced Technology (SIAT), Chinese Academy of Sciences.

### 2.2 Participants

Sixty-six healthy Chinese volunteers (17 females; age: M = 23.34 years, SD = 2.01) participated in this study. All participants reported normal or corrected-to-normal vision and normal color vision. All were right-handed, naïve to the purpose of the study, and provided informed consent before the experiments.

Participants completed the Chinese version of the State-Trait Anxiety Inventory (STAI; Spielberger et al., 1983) after EEG collection. The trait-anxiety scores ranged from 21 to 67 (mean = 41.77, SD = 11.16). Participants’ trait-anxiety score above 50 was defined as high anxiety (n = 18), and scores below 35 as low anxiety (n = 20). The specific STAI cutoffs were chosen according to previous ERP studies of anxiety (Eldar et al., 2010; Gaspar & McDonald, 2018).

### 2.3 Stimuli and Procedures

Stimuli were presented on a 100-Hz LCD monitor with a black background that was viewed from a distance of 60 cm. Figure 1 shows a schematic of the stimuli. A white fixation cross was continuously visible at the center of the screen during the experiment. Each search display contained six shapes distributed at equal distances around a virtual circle with a radius of 7.5° and two of the shapes located on the vertical centerline. The shapes were a diamond (0.9° by 0.9°) and a circle (1.1° diameter) colored either red or green. Each shape contained a gray line tilted either 45° or 135°. The target was designated as the unique geometrical shape among the six items, i.e., a diamond or a circle, which switched randomly between trials. A color-singleton distractor (red or green) appeared during 2/3 of trials (distractor-present trials), in which one of the non-target items had a different color from others. During the remaining 1/3 of the trials (distractor-absent trials), all items were of the same color. The target could be red or green. Target and distractor locations varied to produce the following six types of configurations: lateral target/no distractor (2/9), centerline target/no distractor (1/9), lateral target/centerline distractor (1/6), centerline target/lateral distractor (1/6), lateral target/ipsilateral distractor (1/6), and lateral target/contralateral distractor (1/6).

**Figure. 1.**
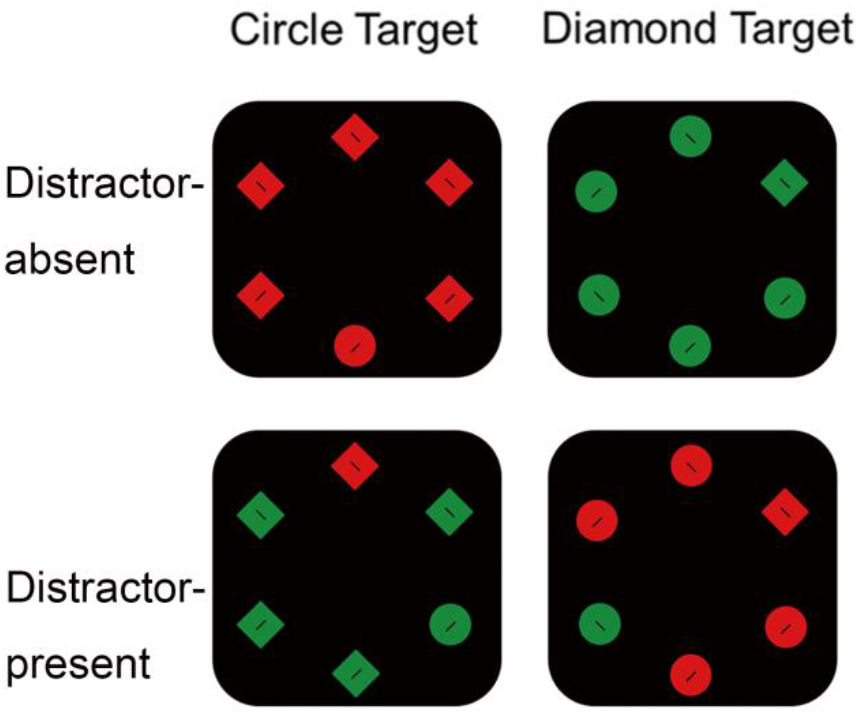
Stimulus examples. Participants were required to report the tilt of a gray line (45° or 135°) inside the unique target shape (circle or diamond). A color distractor appeared randomly in 1/3 of the trials.

On each trial, a search display was presented following an 800- to 1200-ms central fixation. Participants were instructed to maintain central fixation and report the orientation of the tilted line within the target as quickly as possible by pressing the left or right buttons. The search array was presented for 200 ms and response was required within a 2-s time limit after the onset of the search array. Participants were given auditory feedback if the response was incorrect. Participants were informed about the possibility of a color distractor and were asked to ignore it. All the conditions were presented randomly in 1512 trials.

### 2.4 Behavioral measures

Response times (RTs) were analyzed for the distractor-present and -absent conditions. Correct trials with RT between 200 and 2000 ms were analyzed. We exclude the trials in which participants respond too quickly or slowly. In this study, the average RT was 857 ± 78 ms. The specific cut-offs (200 ms and 2000 ms) refer to previous studies using similar stimuli and the average RTs in these studies were also between 800 and 1000 ms (Burra & Kerzel, 2014; McDonald et al., 2012). The interference effect was calculated by the RT difference between the distractor-present and distractor-absent conditions.

### 2.5 EEG recording and analysis

EEG data was acquired from 128 channels (Hydro Cel Geodesic Sensor Net; Electrical Geodesics, Inc., Eugene, OR) with a sampling rate of 1000 Hz (reference: Cz, impedance < 50 kΩ). Offline EEG processing and analyses were performed in MATLAB using the EEGLAB toolbox. EEG signals were resampled offline to 250 Hz, filtered with filter cut-offs of 0.1 Hz and 40 Hz, and re-referenced to the average earlobes (the average of the left and right mastoid channels). Then independent component analysis (ICA) was performed using EEGLAB’s BINICA routine using all electrodes, and ICA components associated with eye blinks, horizontal eye movements and heartbeats were visually identified and removed according to their spatial, spectral, and temporal properties. Epochs extended from 200 ms before stimulus onset to 500 ms after stimulus onset, and a 100-ms pre-stimulus window was used for baseline correction. To further remove the horizontal eye movements in the data, we excluded blinks and vertical eye movements (Fp1/Fp2 exceeding ± 70 V), horizontal eye movements (F9/F10 channels exceeding ± 30 V). An average of 4.2 % of trials were rejected for all participants.

The N2pc and Pd components were measured as contralateral-minus-ipsilateral (contra-minus-ipsi) difference waves. Distractor-evoked ERPs were extracted from the trials of centerline target/lateral distractor (Distractor), and target-evoked N2pc component was extracted from the trials of centerline distractor/lateral target (Tar-Distra-Pre) or no distractor/lateral target (Tar-Distra-Abs). We measured the N2pc and Pd components of the mean waveforms recorded at several electrode sites around PO7 and PO8 (marked by white dots in Fig. 2B), where these components showed large amplitude. The amplitudes were calculated as the average value during the time window 20 ms before and after the peak of averaged N2pc and Pd for all participants (Gaspar & McDonald, 2018; Luck, 2014). The amplitudes and peak latencies of ERP components were analyzed. The peak latencies of N2pc and Pd were measured for each participant during the time window of 150-400 ms, which is the classic time window of N2pc and Pd (Gaspar & McDonald, 2014; Hu et al., 2019).

**Figure. 2.**
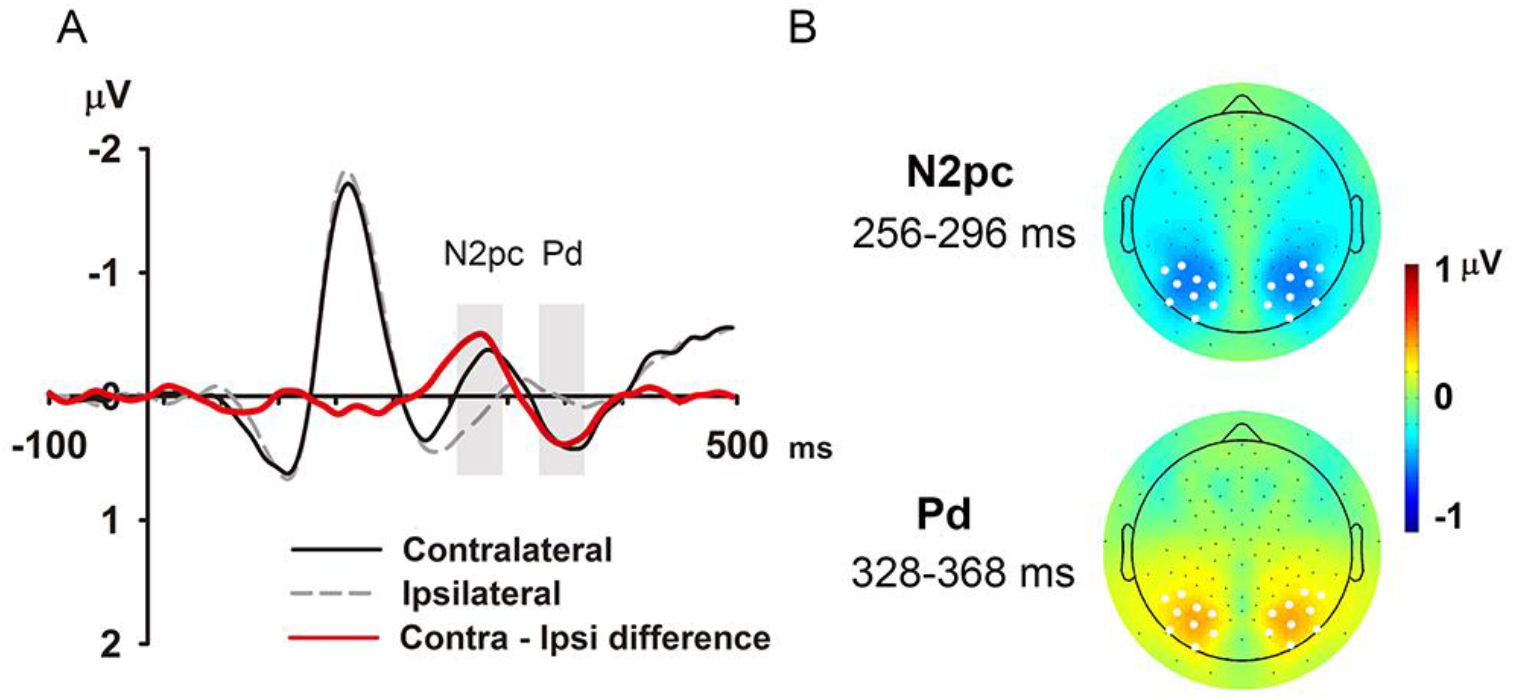
Distractor-evoked ERPs for all participants. (A) Average distractor-evoked ERPs from all participants (n = 66). ERP waveforms recorded contralaterally and ipsilaterally to the color singleton distractor are plotted separately in solid black line and dotted gray line. Contralateral-minus-ipsilateral (contra-ipsi) difference waveforms were shown in the red line. The two shaded boxes represent the time window of the distractor-evoked N2pc (256-296 ms) and Pd (328-368 ms). (B) Topographic maps of N2pc and Pd. The N2pc and Pd components were mainly distributed over posterior areas. The electrodes where we measured N2pc and Pd were marked by white dots.

## 3. Results

### 3.1 Behavioral results

The mean RT of participants was significantly longer in distractor-present trials (886 ms) than in distractor-absent trials (813 ms, *t*(66) = 20.210, *p* < 0.001, Cohen’s d = 2.488), suggesting a strong interference effect of the distractors. However, there was no significant correlation between the behavioral interference effect and the levels of trait anxiety (r = 0.057, *p* = 0.651; BF_10_ = 0.107). The behavioral interference effect (i.e., the RT difference between distractor-present and distractor-absent trials) of the high-anxiety group was numerically greater than that of the low-anxiety group (77 ± 6 ms vs. 73 ± 8 ms), but the difference between the two groups failed to reach statistical significance (t(36) = 0.396, *p* = 0.694, and Cohen’s d = 0.129; BF_10_ = 2.241).

### 3.2 Distractor evoked both N2pc and Pd

To study the attentional processing of the distractor, we analyzed N2pc and Pd components elicited by the distracting color singleton. For all participants, the lateral distractor elicited significant N2pc (256-296 ms) after stimulus presentation (contra-minus-ipsi: −0.431 ± 0.058 μV (mean ± S.E.), *t*(65) = −7.362, *p* < 0.001, and Cohen’s d = −0.906; BF_10_ > 1000; Fig. 2A, red solid line) and significant Pd (328-368 ms) after stimulus presentation (contra-minus-ipsi: 0.314 ± 0.055 μV, *t*(65) = 5.697, *p* < 0.001, and Cohen’s d = 0.701; BF_10_ > 1000; Fig. 2A, red solid line). The results revealed that the distractor captured attention first and then was suppressed quickly, in line with previous studies (Burra & Kerzel, 2013; Hilimire & Corballis, 2014; Hilimire et al., 2011; Hu et al., 2019).

### 3.3 Distractor-evoked Pd amplitudes were negatively correlated with the level of trait anxiety

Pearson correlation analyses revealed a significant negative correlation between Pd amplitudes (328-368 ms) and trait-anxiety scores (r = −0.327, p = 0.007; BF_10_ = 3.441; Fig. 3A), i.e., Pd amplitude decreased with the increasing level of trait anxiety. In contrast, there was no significant correlation between N2pc amplitudes (238-306 ms) and trait-anxiety scores (r = − 0.002, *p* = 0.986; BF_10_ = 0.097; Fig. 3B). Besides, analysis of statistical difference between the two correlations showed that the correlation between Pd amplitudes and anxiety scores was significantly higher than that between N2pc amplitudes and anxiety scores (z = −1.886, *p* = 0.03, refer to the method in Eid et al., 2011). The correlation results showed that the amplitudes of the Pd, but not N2pc, were correlated with trait-anxiety scores, which suggests that the higher the level of anxiety, the worse the ability of attentional inhibition. We didn’t find significant correlation between trait-anxiety scores and peak latencies of distractor-evoked N2pc (286 ± 5 ms, r = 0.161, *p* = 0.194; BF_10_ = 0.222) or Pd (301 ± 10 ms, r = −0.212, *p* = 0.09; BF_10_ = 0.414).

**Figure. 3.**
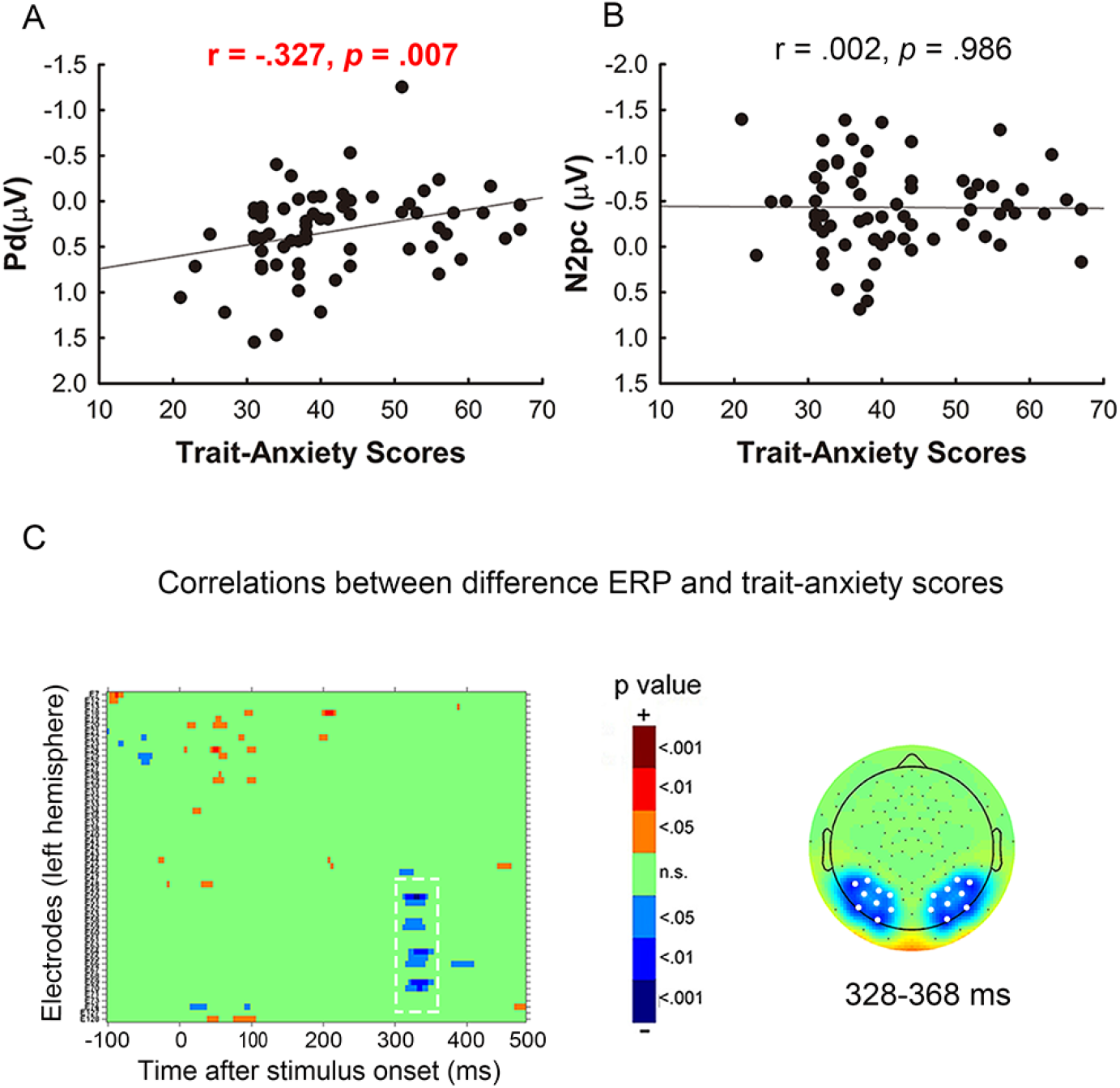
Distractor-evoked ERPs and their correlations with trait-anxiety scores. (A) Correlation between trait-anxiety scores and N2pc amplitudes for all participants. (B) Correlation between trait-anxiety scores and Pd amplitudes for all participants. (C) Statistical significance of correlations between distractor-evoked contralateral-minus-ipsilateral difference ERP and trait-anxiety scores at each electrode site on the left hemisphere (similar results for the right hemisphere). Pearson correlations were carried out with a sliding window of 20 ms and a step of 4 ms. Different colors were used to indicate the significance levels (refer to the color bar). Topographic map of p values of correlations between the Pd interval (328-368 ms) and trait-anxiety scores in the right panel. The electrodes where we measured Pd were marked by white dots.

Further, we examined the correlations between amplitudes of distractor-evoked difference ERP and trait-anxiety scores at each electrode site in the interval of −100 ms to 500 ms (Fig. 3C, left panel). We found that the time range of significant correlations on the posterior electrodes fit well with the Pd time window (about 300-380 ms). And a p-value map of correlations during the Pd time window also showed that significant correlations were concentrated on the posterior electrodes (Fig. 3C, right panel). In sum, the correlation results from 0 to 500 ms on the whole brain suggest that only the Pd component was significantly correlated with trait anxiety scores.

### 3.4 Peak latency of target-evoked N2pc was positively correlated with the level of trait anxiety

We also analyzed the target-evoked ERP. The lateral target elicited significant N2pc (260-300 ms) in both distractor-absent (Tar-Distra-Abs: −0.567 ± 0.077 μV, *t*(65) = −7.352, *p* < 0.001, and Cohen’s d = −0.905; BF_10_ >1000; Fig. 4A, red solid line) and distractor-present conditions (Tar-Distra-Pre: −0.183 ± 0.044 μV, *t*(65) = −4.111, *p* < 0.001, and Cohen’s d = −0.506; BF_10_ = 273.776; Fig. 4A, blue solid line). We didn’t find significant correlation between trait-anxiety scores and amplitudes of target-evoked N2pc in either distractor-absent condition (r = 0.166, *p* = 0.180; BF_10_ = 0.234) or distractor-present condition (r = 0.054, *p* = 0.662; BF_10_ = 0.106). However, there was a significant correlation between trait-anxiety scores and peak latencies of target-evoked N2pc in distractor-absent condition (296 ± 5 ms, r = 0.358, *p* = 0.003; BF_10_ = 7.355; Fig. 4C), while no significant correlation in distractor-absent condition (290 ± 10 ms, r = 0.028, *p* = 0.822; BF_10_ = 0.099). Fig. 4D showed p values of correlations between trait-anxiety scores and target-evoked negative peak latency from 150 to 400 ms at each electrode site in distractor-absent condition.

**Figure. 4.**
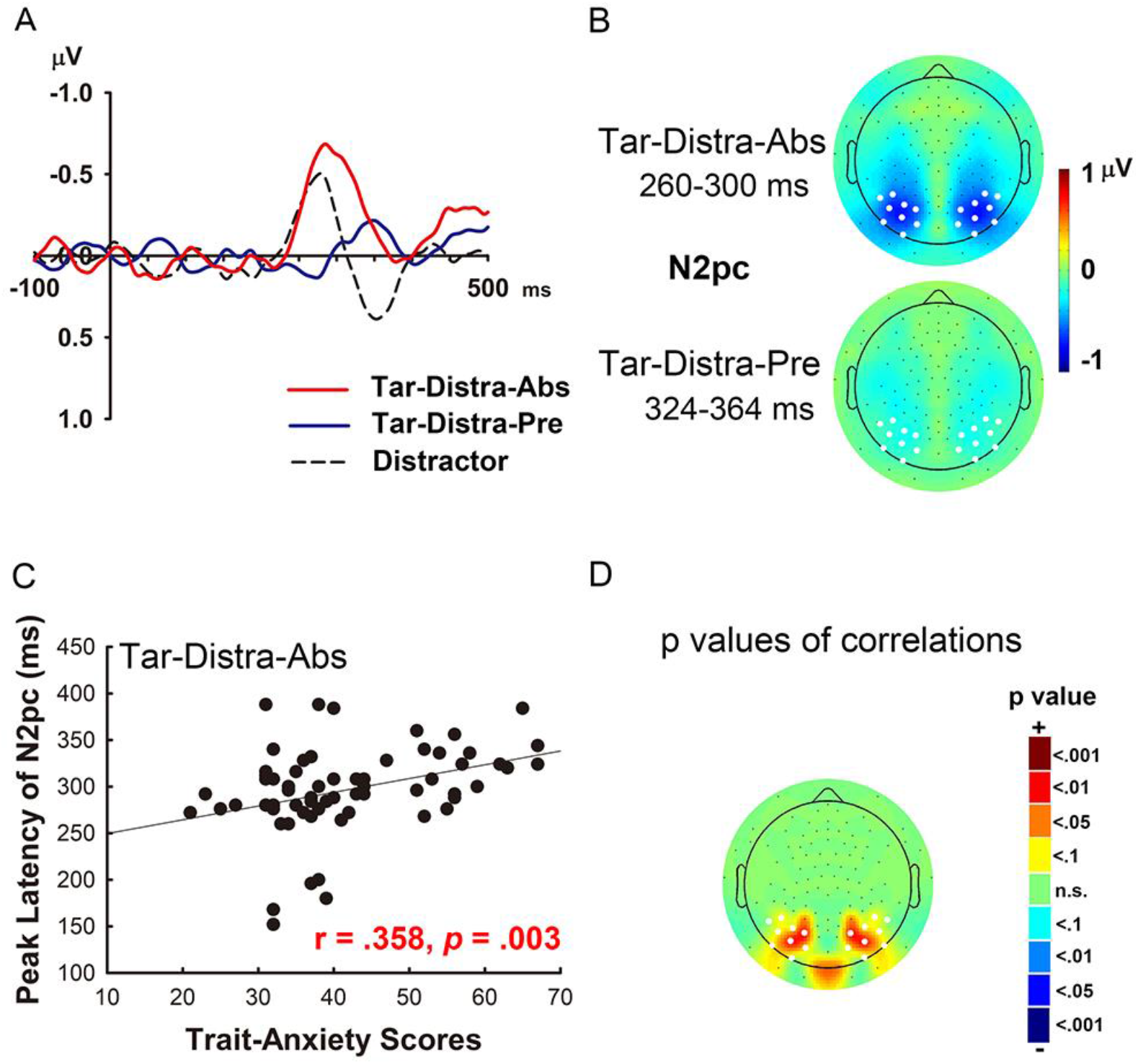
Target-evoked ERPs and their correlations with trait-anxiety scores. (A) Contralateral-minus-ipsilateral difference waveforms of the target in distractor-absent (Tar-Distra-Abs, red line) condition and distractor-present condition (Tar-Distra-Pre, blue line), and of the distractor (black dot line). (B) Topographic maps of the target-evoked N2pc in distractor-absent and distractor-present conditions. The electrodes where we measured the N2pc were marked by white dots. (C) Correlation between trait-anxiety scores and peak latencies of the N2pc for all participants. The peak latency was the latency of the smallest value during 100-400 ms. (D) Topographic map of p-values of the correlations between peak latencies of the N2pc and trait-anxiety scores. The electrodes where we measured the Pd were marked by white dots.

### 3.5 Comparisons between high and low anxiety groups

Most of previous ERP studies on anxiety have investigated differences between high and low anxiety groups (e.g., Eldar et al., 2010; Gaspar & McDonald, 2018). Therefore, we analyzed the data from participants whose trait-anxiety scores were above 50 (n = 18, high-anxiety group) or below 35 (n = 20, low-anxiety group) to directly compare ERP differences between high and low anxiety groups. Based on the findings of correlation analysis, we analyzed the differences between the two groups in these anxiety-related indicators.

As shown in Fig. 5A, C and E, the Pd amplitude difference between the high- and low-anxiety groups was significant (*t*(36) = 2.712, *p* = 0.010, and Cohen’s *d* = 0.881; BF_10_ = 6.810; Fig. 5A): a significant Pd was found in the low-anxiety group (0.528 ± 0.058 μV, *t*(19) = 5.047; p < 0.001, Cohen’s *d* = 1.129; BF_10_ = 432.046; Fig. 5A, red solid line) while there was no significant Pd component in the high-anxiety group (0.126 ± 0.054 μV, *t*(17) = 1.212; *p* = 0.242, Cohen’s *d* = 0.286; BF_10_ = 1.620; Fig. 5A, blue solid line). In contrast, there was no significant difference in N2pc amplitudes between the two groups (−0.460 ± 0.058 μV vs. −0.490 ± 0.043 μV, *t*(36) = 0.222; *p* = 0.825, and Cohen’s *d* = 0.072; BF_10_ = 2.345; Fig. 5A), suggesting that the distractor captured attention to a similar degree for the two groups. The results suggest that the difference between the high- and low-anxiety groups is mainly reflected in the top-down inhibition process of attention, not the bottom-up attentional process.

**Figure 5.**
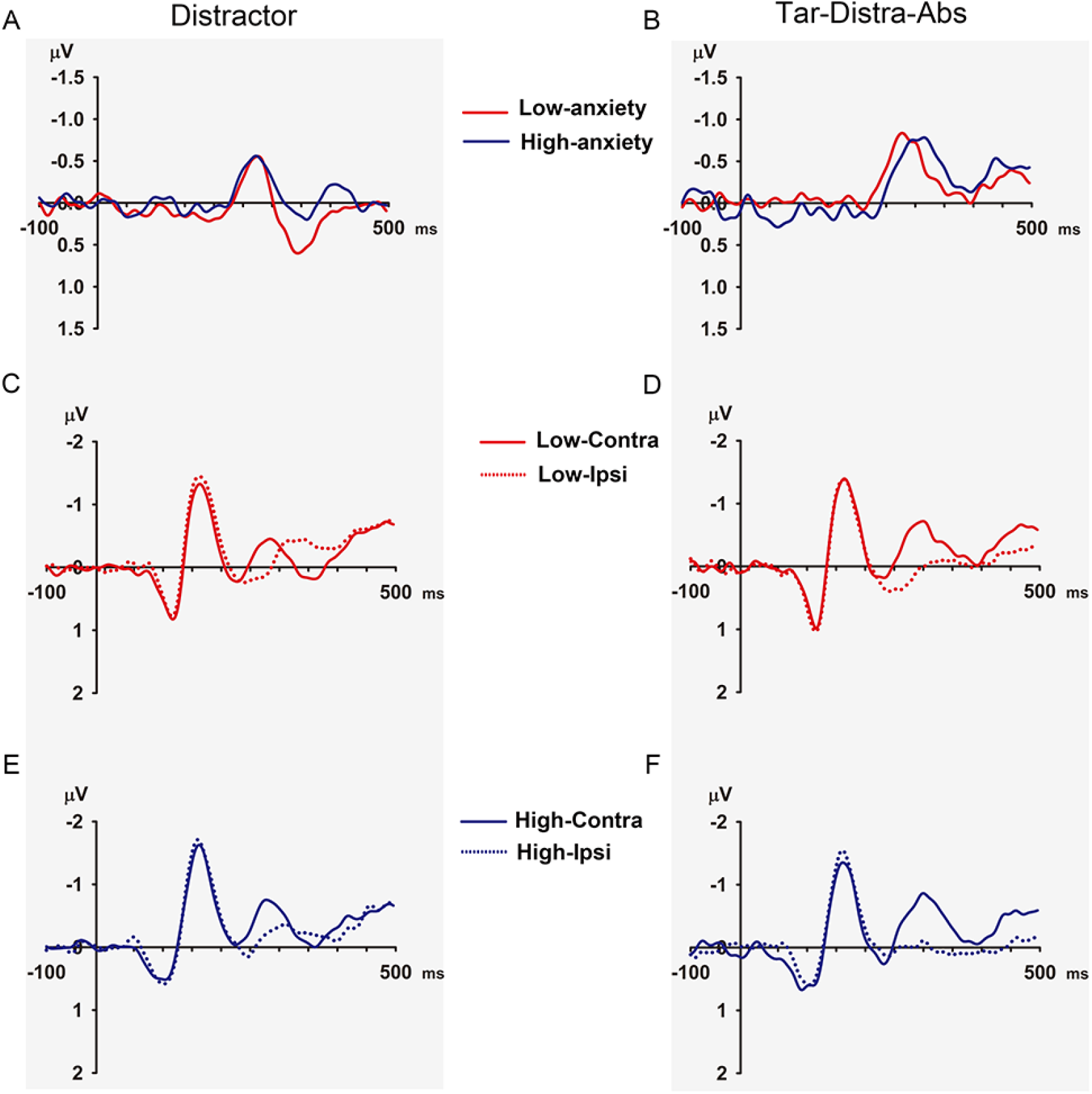
Significant ERPs difference between the high- and low-anxiety groups. Contralateral-minus-ipsilateral difference waveforms of the distractor (A) and the target (B) in distractor-absent condition for the low-anxiety (n = 20) and high-anxiety groups (n = 18). The results showed that the Pd was non-significant in the high-anxiety group while it was robust in the low-anxiety group. Peak latency of target-evoked N2pc in distractor-absent trials was significantly delayed for the high-anxiety group compared with that for the low-anxiety group. Average ERP signals triggered by the lateral salient distractor for the low-anxiety (C) and high-anxiety (E) groups, and by the lateral target in distractor-absent condition for the low-anxiety (D) and high-anxiety (F) groups.

Fig. 5B, D, and F showed target-evoked ERPs in the distractor-absent condition for the high- and low-anxiety groups. We found that the peak latency of N2pc was significantly different between the two groups (*t*(36) = −2.786, *p* = 0.008, and Cohen’s *d* = −0.905; BF_10_ = 7.814; Fig. 5B): an earlier N2pc was found in the low-anxiety group (282.200 ± 6.294 ms) and a later one in the high-anxiety group (320.889 ± 3.786 ms). There was no significant difference in the amplitude of N2pc component (256-296 ms) between the two groups (*t*(36) = −1.057, *p* = 0.297, and Cohen’s *d* = −0.343; BF_10_ = 1.503). The results suggest that high-anxiety individuals take a longer time to attend the target compared to low-anxiety individuals.

### 3.6 Internal consistency reliability of behavioral and ERP measures

Internal consistency reliability of behavioral and ERP measures was examined with a split-half approach, where the correlation between averages of odd- and even-numbered trials was determined and corrected using the Spearman-Brown prophecy formula (Nunnally, Bernstein, & Berge, 1967). The Spearman-Brown corrected correlations of behavioral and ERP measures were all significant (all r > 0.907, all *ps* < 0.001). We also examined the internal consistency reliability of trait anxiety report. Cronbach’s alpha for the STAI (trait anxiety scores) was 0.922. These results indicated that internal consistency reliability across all analytic strategies was acceptable to good.

## 4. Discussion

The present study investigates the neural mechanism underlying anxiety-related attentional impairment by recording high-resolution EEG in a visual search task with unpredictable targets and distractors. The relationships of trait-anxiety scores and ERP components N2pc and Pd were concerned. The N2pc is the marker of attentional selection (Eimer, 1996; Hickey et al., 2006), and the Pd is the marker of attentional inhibition (Hickey et al., 2009; Sawaki & Luck, 2010). We found a negative correlation between trait-anxiety scores and distractor-evoked Pd amplitudes, that is, the higher the level of anxiety, the worse the attentional inhibition. In contrast, there was no significant correlation between anxiety scores and distractor-evoked N2pc amplitudes. Further, we compared the distractor-evoked ERPs of the high- and low-anxiety groups. The results showed that the Pd was non-significant in the high-anxiety group while it was robust in the low-anxiety group. In contrast, the N2pc did not differ between the two groups. In addition, we also found a positive correlation between trait-anxiety scores and the peak latencies of target-evoked N2pc when the distractor was absent, suggesting that higher-level anxiety may induce delayed attentional selection of the target. In sum, the present findings suggest that an individual’s ability in attentional inhibition of the distractor and attentional selection of the target correlates with trait anxiety level.

A novel and crucial finding of our study is that we found direct neurophysiological evidence for impairments in attentional inhibition in anxiety. That is, the amplitude of distractor-evoked Pd decreases with trait-anxiety levels, which means that individuals with higher-level anxiety have more difficulty inhibiting task-irrelevant information. Although behavioral studies and anxiety-related theories suggest anxiety disrupts attentional inhibition, the neural correlates are far less understood (Berggren & Derakshan, 2013). Our findings suggest that anxiety disrupts the ability to inhibit task-irrelevant information and the degree of anxiety-related impairment in the attentional inhibition could be quantitatively assessed using the Pd component.

We observed distractor-evoked N2pc and the following Pd in a visual search task with unpredictable targets and distractors which changed shape and color from trial to trial. The finding is consistent with previous studies (Burra & Kerzel, 2013; Hilimire & Corballis, 2014; Hilimire et al., 2011). If a target or a distractor has a fixed feature (i.e., it is predictable), the distractor-evoked N2pc would decrease or disappear (Feldmann-Wüstefeld & Anna Schubö, 2016; Gaspar & McDonald, 2014; Hilimire & Corballis, 2014; Jannati et al., 2013). A recent study found that the predictability of a distractor could also reduce the amplitude of Pd (van Moorselaar et al., 2020). On the contrary, unpredictability could bring robust the N2pc and the Pd. Thus in our task, we observed distractor processing, including attentional capture and inhibition indexed by the N2pc and Pd.

Our finding of a difference in distractor-evoked Pd instead of N2pc between the high- and low-anxiety groups seems to be inconsistent with a previous study (Gaspar & McDonald, 2018). However, we think that the two findings are not necessarily contradictory. In their study, the target and distractor were predictable. They found that the distractor-evoked N2pc was not observed in the low-anxiety group while the N2pc appeared in the high-anxiety group. Under a paradigm with predictable stimuli, the main difference in attentional processing induced by anxiety lies in the process of bottom-up attention. In our study, robust N2pc and Pd components were observed in both high- and low-anxiety groups as the target and distractor were both unpredictable. The unpredictability of stimuli accounts for our findings that attention capture by the distractor was hard to suppress proactively for both groups (i.e., evoking the N2pc) and then a strong subsequent attentional inhibition was required (i.e., evoking the Pd). Therefore, anxiety-related impairment in attention mainly exhibits in the process of top-down inhibition, thus the Pd disappeared in the high-anxiety group. We speculate that both bottom-up and top-down processes play an important role in the anxiety-related interference effect, with the weight of the processes varying under different requirements of attentional inhibition.

In terms of behavioral performance, anxiety levels did not seem to significantly affect behavioral interference effect. Nevertheless, the trend toward greater interference in the high anxiety group is consistent with the present ERP finding of impaired Pd component in the high anxiety group. In fact, difference in statistical significance between behavioral performance and brain activities was also reported in previous studies (e.g., Harris et al., 2020; Gasper et al., 2018). One possible reason is that behavioral results are the output of complex multiple processes, whereas EEG measures reflect more direct processing.

Another interesting finding is that attentional selection of the target slows down in individuals with high-level anxiety, reflected in a delayed latency of target-evoked N2pc, which is considered to represent the speed of attentional selection (Foster et al., 2020; Bachman et al., 2020). Findings from previous studies suggest that it is not consistent in anxiety-related impairments of N2pc evoked by a non-emotional target. Some evidence shows that the amplitude of neutral-target-evoked N2pc decreased with the increase of anxiety level (Moran & Moser, 2015), while some research did not find a significant change in the N2pc (Gaspar & McDonald, 2018). Here we find that the peak latencies of N2pc positively correlates with trait-anxiety levels. A trend of delayed target-evoked N2pc in high anxiety group was also reported by Gaspar & McDonald (2018) even though the result failed to reach significance. The findings suggest that anxiety affects the efficiency of top-down attentional selection of the target.

The findings of the present study contribute to anxiety-related theories in two ways. First, top-down attentional inhibition is a core function of attentional control, which is highlighted in most cognitive models of anxiety. In EEG studies, the Pd component is considered a direct marker of top-down inhibition. For the first time, we find that the Pd reduces as a function of trait anxiety. The correlation between Pd amplitudes and levels of anxiety provides quantitative data for the updating of anxiety-related theories. Second, increasing behavioral evidence shows that anxious individuals tend to be distracted by irrelevant stimulation not only for threat-related stimuli but also for non-emotional neutral stimuli anxiety (Moser et al., 2012). Based on these findings, some anxiety-related theories propose that anxiety is linked to a general impairment of the attentional system (Eysenck & Derakshan, 2011). Our EEG findings with non-emotional stimuli suggest that the anxiety-related deficit is a general attentional dysfunction, specifically in attentional inhibition. In addition, our new finding of anxiety delays the attentional selection of a non-emotional target have implications for updating theories about anxiety.

## 5. Conclusions

In summary, the present ERP study provides direct neurophysiological evidence that anxiety is associated with a general impairment in attentional control, especially in attentional inhibition. The present finding provides access to the mechanisms underlying the anxiety-related impairments in attentional control: high anxiety disrupts attentional inhibition of the distractor (indexed by the Pd) and the efficiency of top-down attention selection of the target (indexed by the N2pc). We believe that the N2pc and Pd components may be potential neurophysiological indicators for the degree of attentional impairment in anxiety disorders. Future work will recruit more participants, including anxiety patients, to investigate anxiety-related attention disorders.

## Credit authorship contribution statement

**Liping Hu**: Conceptualization, Software, Formal analysis, Writing- Original draft. **Huikang Zhang**: Investigation, Data curation. **Hongsi Tang**: Investigation. **Lin Shen**: Investigation, Validation. **Rui Wu**: Investigation. **Yan Huang**: Conceptualization, Methodology, Formal analysis, Writing- Reviewing and Editing, Supervision, Resources.

## Acknowledgments

The study was funded by Guangdong Provincial Key Laboratory of Brain Connectome and Behavior (2017B030301017), CAS Key Laboratory of Brain Connectome and Manipulation (2019DP173024), Science and Technology Program of Guangdong (2018B030334001), the National Natural Science Key Foundation of China (81830040), Jiangsu Provincial Medical Outstanding Talent (JCRCA2016006).

## Data statement

Raw data used for analyses presented within this article will be made available upon request. If you would like to access the data, please send an e-mail to Yan Huang at the following e-mail address.

## References

Bachman, M. D., Wang, L. L., Gamble, M. L. & Woldorff, M. G. (2020). Physical Salience and Value-Driven Salience Operate through Different Neural Mechanisms to Enhance Attentional Selection. Journal of Neuroscience, 40, 5455–5464. https://doi.org/10.1523/JNEUROSCI.1198-19.2020

Bar-Haim, Y., Lamy, D., Pergamin, L., Bakermans-Kranenburg, M.J., & van Ijzendoorn, M. H. (2007). Threat-related attentional bias in anxious and nonanxious individuals: A meta-analytic study. Psychological Bulletin, 133, 1–24. https://doi.org/10.1037/0033-2909.133.1.1

Basten, U., Stelzel, C., & Fiebach, C. J. (2011). Trait anxiety modulates the neural efficiency of inhibitory control. Journal of Cognitive Neuroscience, 23, 3132–3145. https://doi.org/10.1162/jocn_a_00003

Berggren, N., & Derakshan, N. (2013). Attentional control deficits in trait anxiety: why you see them and why you don’t. Biological Psychology, 92, 440–446. https://doi.org/10.1016/j.biopsycho.2012.03.007

Berggren, N., & Derakshan, N. (2014). Inhibitory deficits in trait anxiety: Increased stimulus-based or response-based interference? Psychonomic Bulletin & Review, 21, 1339–1345. https://doi.org/10.3758/s13423-014-0611-8

Bishop, S. J. (2009). Trait anxiety and impoverished prefrontal control of attention. Nature Neuroscience, 12, 92–98. https://doi.org/10.1038/nn.2242

Burra, N., & Kerzel, D. (2013). Attentional capture during visual search is attenuated by target predictability: Evidence from the N2pc, Pd, and topographic segmentation. Psychophysiology, 50, 422–430. https://doi.org/10.1111/psyp.12019

Dennis, T. A., & Chen, C. C. (2009). Trait anxiety and conflict monitoring following threat: an ERP study. Psychophysiology, 46, 122–131. https://doi.org/10.1111/j.1469-8986.2008.00758.x

Eid, M., Gollwitzer, M., & Schmitt, M. (2011). Statistik und Forschungsmethoden Lehrbuch. Weinheim: Beltz.

Eimer, M. (1996). The N2pc component as an indicator of attentional selectivity. Electroencephalography and Clinical Neurophysiology, 99, 225–234. https://doi.org/10.1016/0013-4694(96)95711-9

Eldar, S., Yankelevitch, R., Lamy, D., & Bar-Haim, Y. (2010). Enhanced neural reactivity and selective attention to threat in anxiety. Biological Psychology, 85, 252–257. https://doi.org/10.1016/j.biopsycho.2010.07.010

Eysenck, M. W., & Derakshan, N. (2011). New perspectives in attentional control theory. Personality and Individual Differences, 50, 955–960. https://doi.org/10.1037/1528-3542.7.2.336

Eysenck, M. W., Derakshan, N., Santos, R., & Calvo, M. G. (2007). Anxiety and cognitive performance: Attentional control theory. Emotion, 7, 336–353. https://doi.org/10.1037/1528-3542.7.2.336

Feldmann-Wüstefeld, T., & Schubö, A. (2016). Intertrial priming due to distractor repetition is eliminated in homogeneous contexts. Attention, Perception, and Psychophysics, 78, 1935–1947. https://doi.org/10.3758/s13414-016-1115-6

Gaspar, J. M., & McDonald, J. J. (2014). Suppression of salient objects prevents distraction in visual search. The Journal of Neuroscience, 34, 5658–5666. https://doi.org/10.1523/JNEUROSCI.4161-13.2014

Foster, J. J., Bsales, E. M. & Awh, E. (2020). Covert Spatial Attention Speeds Target Individuation. Journal of Neuroscience, 40, 2717–2726. https://doi.org/10.1523/JNEUROSCI.2962-19.2020

Gaspar, J. M., & McDonald, J. J. (2018). High Level of Trait Anxiety Leads to Salience-Driven Distraction and Compensation. Psychological Science, 29, 2020–2030. https://doi.org/10.1037/emo0000506

Gaspelin, N., & Luck, S. J. (2017). The role of inhibition in avoiding distraction by salient stimuli. Trends in Cognitive Sciences, 22, 79–92. https://doi.org/10.1016/j.tics.2017.11.001

Gutierrez, M., & Berggren, N. (2020). Anticipation of aversive threat potentiates task-irrelevant attentional capture. Cognition and Emotion, 34, 1036–1043. https://doi.org/10.1080/02699931.2019.1706448

Harris, A., Jacoby, O., Remington, R., Becker, S., & Mattingley, J. (2020) Behavioral and electrophysiological evidence for a dissociation between working memory capacity and feature-based attention, Cortex, 129, 158–174. https://doi.org/10.1016/j.cortex.2020.04.009

Hickey, C., Di Lollo, V., & McDonald, J. J. (2009). Electrophysiological indices of target and distractor processing in visual search. Journal of Cognitive Neuroscience, 21, 760–775. https://doi.org/10.1162/jocn.2009.21039

Hickey, C., McDonald, J. J., & Theeuwes, J. (2006). Electrophysiological evidence of the capture of visual attention. Journal of Cognitive Neuroscience, 18, 604–613. https://doi.org/10.1162/jocn.2006.18.4.604

Hilimire, M. R., & Corballis, P. M. (2014). Event-related potentials reveal the effect of prior knowledge on competition for representation and attentional capture. Psychophysiology, 51, 22–35. https://doi.org/10.1111/psyp.12154

Hilimire, M. R., Mounts, J. R., Parks, N. A., & Corballis, P. M. (2011). Dynamics of target and distractor processing in visual search: Evidence from event-related brain potentials. Neuroscience Letters, 495, 196–200. https://doi.org/10.1016/j.neulet.2011.03.064

Hu, L., Ding, Y., & Qu, Z. (2019). Perceptual learning induces active suppression of physically nonsalient shapes. Psychophysiology, 56, e13393. https://doi.org/10.1111/psyp.13393

Jannati, A., Gaspar, J. M., & McDonald, J. J. (2013). Tracking target and distractor processing in fixed- feature visual search: Evidence from human electrophysiology. Journal of Experimental Psychology: Human Perception and Performance, 39, 1713–1730. https://doi.org/10.1037/a0032251

Kalanthroff, E., Henik, A., Derakshan, N., & Usher, M. (2016). Anxiety, emotional distraction, and attentional control in the Stroop task. Emotion 16, 293–300. https://doi.org/10.1037/emo0000129

Luck, S. J., & Hillyard, S. A. (1994). Electrophysiological correlates of feature analysis during visual search. Psychophysiology, 31, 291–308. https://doi.org/10.1111/j.1469-8986.1994.tb02218.x

McTeague, L. M., Shumen, J. R., Wieser, M. J., Lang, P. J., & Keil, A. (2011). Social vision: Sustained perceptual enhancement of affective facial cues in social anxiety. NeuroImage, 54, 1615–1624. https://doi.org/10.1016/j.biopsych.2017.10.004

Mogg, K., & Bradley B. P. (2018). Anxiety and Threat-Related Attention: Cognitive-Motivational Framework and Treatment. Trends in Cognitive Sciences, 22, 225–240. https://doi.org/10.1016/j.tics.2018.01.001

van Moorselaar, D., Lampers, E., Cordesius, E., & Slagter, H. A. (2020). Neural mechanisms underlying expectation-dependent inhibition of distracting information. Elife, 9, e61048. https://doi.org/10.7554/eLife.61048

Moran, T. P., & Moser, J. S. (2015). The color of anxiety: Neurobehavioral evidence for distraction by perceptually salient stimuli in anxiety. Cognitive, Affective, & Behavioral Neuroscience, 15, 169–179. https://doi.org/10.3758/s13415-014-0314-7

Moser, J. S., Becker, M. W., & Moran, T. P. (2012). Enhanced attentional capture in trait anxiety. Emotion, 12, 213–216. https://doi.org/10.1037/a0026156

Nunnally, J. C., Bernstein, I. H., & Berge, J. M. F. (1967). Psychometric theory (McGraw-Hill Series in Psychology, Vol. 226).

Osinsky, R., Gebhardt, H., Alexander, N., & Hennig, J. (2012). Trait anxiety and the dynamics of attentional control. Biological Psychology, 89, 252–259. https://doi.org/10.1016/j.biopsycho.2011.10.016

Pacheco-Ungietti, A. P., Acosta, A., Callejas, A., & Lupianez, J. (2010). Attention and anxiety: Different attentional functioning under state and trait anxiety. Psychological Science, 21, 298–304. https://doi.org/10.1177/0956797609359624

Sawaki, R., & Luck, S. J. (2010). Capture versus suppression of attention by salient singletons: Electrophysiological evidence for an automatic attend-to-me signal. Attention, Perception, & Psychophysics, 72, 1455–1470. https://doi.org/10.3758/APP.72.6.1455

Spielberger, C. D., Gorsuch, R. L., Lushene, R., Vagg, P. R., & Jacobs, G. A. (1983). Manual for the State-Trait Anxiety Inventory. Sunnyvale, CA: Consulting Psychologists Press

Wang, E., Sun, L., Sun, M., Huang, J., Tao, Y., Zhao, X., Wu, Z., Ding, Y., Newman D. P., Bellgrove M. A., Wang Y. & Song Y. (2016). Attentional Selection and Suppression in Children with Attention-Deficit/Hyperactivity Disorder. Biological Psychiatry, 1, 372–380. http://dx.doi.org/10.1016/j.bpsc.2016.01.004

Wieser, M. J., Pauli, P., & Muhlberger, A. (2009). Probing the attentional control theory in social anxiety: An emotional saccade task. Cognitive, Affective & Behavioral Neuroscience, 9, 314–322. https://doi.org/10.3758/CABN.9.3.314

Wu, J. & Yan J. (2017). Editorial: Stress and Cognition. Frontiers in Psychology, 8, 970. https://doi.org/10.3389/fpsyg.2017.00970

